# Multicellular tumor spheroids- an effective *in vitro* model for understanding drug resistance in head and neck cancer

**DOI:** 10.1101/702316

**Authors:** Mohammad Azharuddin, Karin Roberg, Ashis Kumar Dhara, Mayur Vilas Jain, Padraig D’arcy, Jorma Hinkula, Nigel K H Slater, Hirak K Patra

## Abstract

A hallmark of cancer is the ability to develop resistance against therapeutic agents. Therefore, developing effective *in vitro* strategies to identify drug resistance remains of paramount importance for treatment success. A way cancer cells achieve drug resistance is through the expression of efflux pumps that actively pump drugs out of the cells. To date, several studies have investigated the potential of using 3D multicellular tumor spheroids (MCSs) to assess drug resistance; however, a unified system that uses MCSs to differentiate between multi drug resistant (MDR) and non-MDR cells does not exist. In the present report, we have used MCSs obtained from post-diagnosed, pre-treated (PDPT) patient derived head and neck squamous cancer cells that often become treatment resistant, to develop an integrated approach combining clinical drug response and cytotoxicity screening, real-time drug uptake monitoring, and drug transporter activity assessment using flow cytometry in the presence and absence of their respective specific inhibitors. The present report shows a comparative response to MDR, drug efflux capability, and reactive oxygen species (ROS) activity to assess the resistance profile of PDPT patient-derived MCSs and two-dimensional cultures of the same set of cells. We show that MCSs serve as robust and reliable models for the clinical evaluation of drug resistance. Our proposed strategy can thus have potential clinical applicability for profiling drug resistance in cancers with unknown resistance profiles, which consequentially can indicate benefit from downstream therapy.

## Introduction

Anticancer drug resistance is an unmanageable outcome of a cascade of events that are altered in cancer cells during disease progression. Multidrug resistance (MDR) is one of the mechanisms of anticancer drug resistance, which is described as the resistance to multiple chemotherapeutic drugs with differing structures and functional activities^1,2^. MDR is considered as the major impediment to the success of chemotherapy^3^, and leads to an unprecedented decrease in the survival rate^4^ of cancer patients. The development of MDR occurs at an alarmingly high rate during the treatment phase of various cancers^5^ and the underlying mechanisms of MDR in cancer and subsequent relapse have puzzled researchers worldwide^6^. Only a small subset of tumor cells have been reported to be sufficient for progressive resistance to chemotherapy, leading to the development of MDR in at least 50% of cancer patients^7^. The basic underlying MDR mechanism is associated with 5 events: (i) increased drug efflux, (ii) decreased drug influx, (iii) increased drug metabolism, (iv) increased DNA repair and (v) decreased apoptosis^8,9^. The major players involved in drug efflux related MDR mechanisms are the ATP-binding cassette (ABC) transporter proteins, such as P-glycoprotein (P-gp/MDR1), multidrug resistance-associated protein 1 (MRP-1), and breast cancer resistance protein (BCRP)^10^. Overexpression of P-gp/MDR-1, a membrane-bound active drug efflux pump, appears to be the most prominent contributor to MDR development in cancer cell lines.^11,12^

A direct proportional relationship between elevated ABC transporter levels and MDR progression has been previously reported ^10,13,14^. Presently, well-defined *in vitro* models and assay systems that enable the classification of resistance into MDR and non-MDR categories are limited. First of the two presently employed strategies uses treatment-sensitive *in vitro* cell lines that are exposed to a specific therapeutic anticancer drug until the designated cell line attains a resistance genotype^15^. The second strategy uses a genotype-based assay that focuses on the identification of genetic anomalies arising in the treatment-resistant cell lines^16^. These two tactics have been exploited to integrate numerous MDR pump inhibitors into cancer treatment modalities; however, the outcomes were not sufficiently effective for clinical translation^17^. These strategies have been associated with various discrepancies concerning the differentiation between treatment-sensitive and treatment-resistant cancer cells *in vitro*^10^.

Multicellular tumor spheroids (MCSs) are considered to be the most relevant pre-clinical, high throughput *in vitro* models^18^. MCSs are self-assembled aggregates of cancer cells, which can mimic the complex micro-environmental milieu of the tumor tissue observed *in vivo*^19^. Sutherland’s integration of *in vitro* three-dimensional (3D) culture methods into cancer research nearly four decades ago, triggered an increased interest in the application of MCSs in drug discovery and understanding of the basic biological mechanisms underlying tumor progression and response to treatment^20^. MCSs show an intermediate but clinically relevant complexity between *in vitro* two-dimensional (2D) cell cultures and *in vivo* solid tumors, and they have been assigned a relevant platform for *in vitro* drug screening^21^. They mimic the complex cell-cell adhesion and cell-matrix interactions in solid tumors, which results in gradient generation for nutrients and growth factor signals as observed *in vivo*^19^. In accordance with metabolite gradient and a complex microenvironment, MCSs contain proliferating, quiescent, and necrotic zones, much like the internal milieu of human tumors^22^. In addition, owing to their multicellular nature, MCSs spontaneously develop MDR against many chemotherapeutic drugs^23,24^, thus making them the appropriate model system for the purpose of the present study. Recently, members of our group reported a marked treatment response difference between 2D cell cultures and MCSs of head and neck cancer cells pertaining to epithelial-mesenchymal transition and stem cell characteristics, suggesting that 3D cell cultures are clinically relevant models that are superior to 2D monolayers for the investigation of new therapeutic targets^25^. However, there is no well-defined *in vitro* method or criteria for the identification of the resistance status of cancer cells. Furthermore, integration of these two approaches into translational research is challenging and likely non-implementable in the near future.

In the present study, we describe a fast and robust *in vitro* model and assay system for the profiling of drug resistance status in cancer cells using MCSs obtained from untreated patient-derived PDPT head and neck squamous cancer cells (HNSCCs). This report constitutes a comparative investigation between 2D and MCSs for the assessment of drug resistance profile of the same cell. Our strategy combines drug screening, real time, time-lapse fluorescence microscopy, and flow cytometry for rapid identification of drug resistance status using MCSs, so that a beneficial personalized treatment regimen can be offered to patients. The cell lines used were previously established by the members of our group^25,26^. Briefly, we have investigated the drug response profiles of LK0917 (gingiva), LK0902 (tongue), and LK1108 (hypopharynx) cells to doxorubicin, cisplatin, and methotrexate in 2D and MCSs. In order to establish the drug response profiles for these cell lines, we first investigated their efflux pump activities by assessing the differential uptake of calcein acetoxymethyl ester (calcein-AM), a substrate for the P-gp and MRP1 efflux pumps^27^, using real-time live cell fluorescence imaging^28^. We further studied the reactive oxygen species (ROS) generation in both *in vitro* models using the 2’,7’-dichlorofluorescein diacetate (DCFDA) assay, in order to have a better understanding of the MCS microenvironment of the patient-derived HNSCC cell lines, which we then used for further assessment of MDR status. Finally, we validated our findings with a flow cytometry-based assay for functional detection and profiling of MDR phenotypes in 2D cell cultures and MCSs by assessing calcein-AM uptake in the presence of specific efflux pump inhibitors.

## Materials and methods

### Study design

The schematic representation of the *in vitro* experimental workflow used for determining the MDR profile is provided below (**Scheme 1**). The cell lines LK0912, LK0917, and LK1108 used in our experiments, were established from three different HNSCC patients as reported previously by our team ^25,26^.

**Scheme 1:**
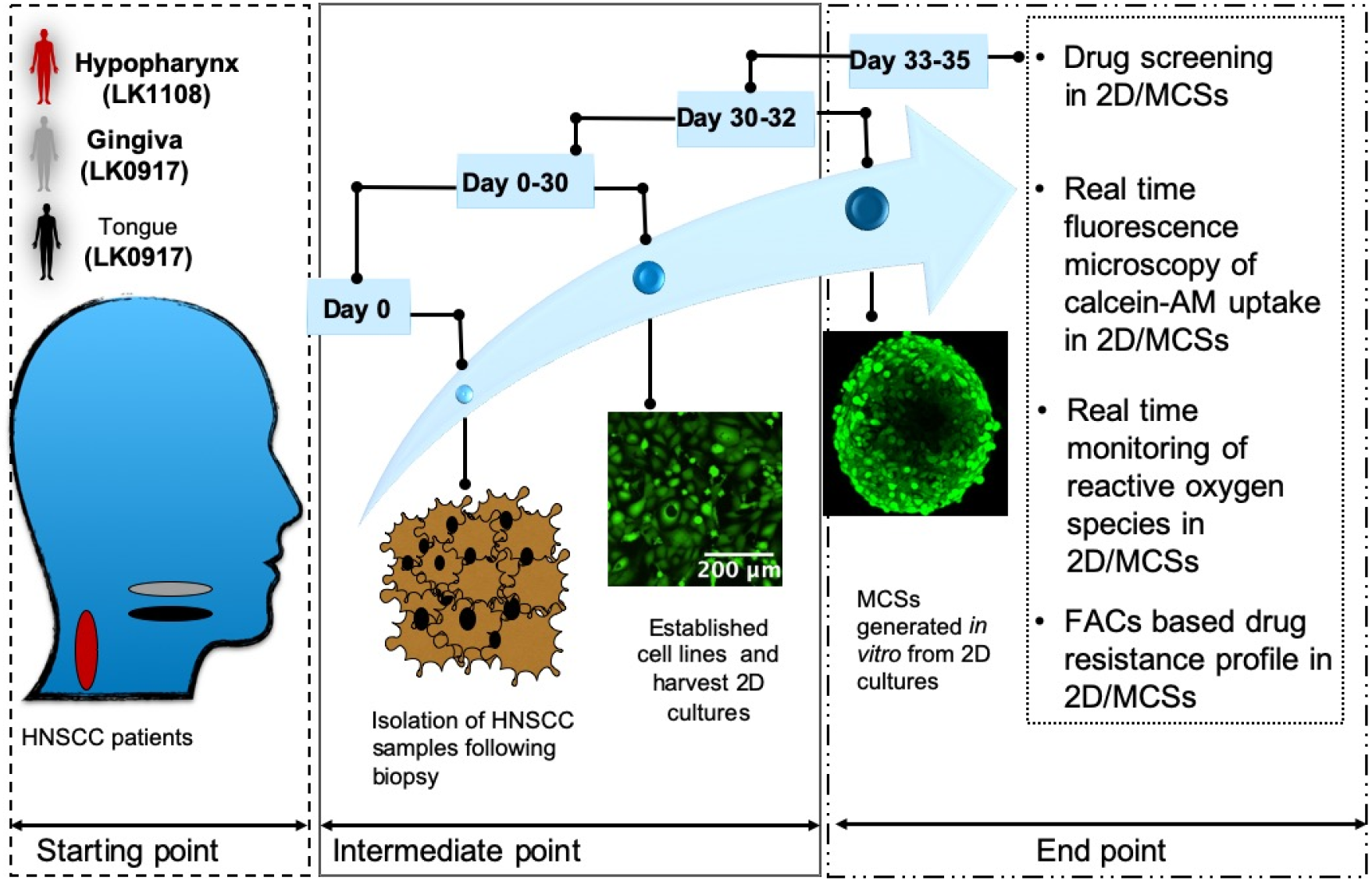
Experimental workflow for MDR screening of cancer cells using MCSs. At the starting point, tumor biopsies from the gingiva, tongue, and hypopharynx of 3 HNSCC patients were obtained. During the intermediate point, cell lines were established from the tumor biopsies and in vitro MCSs cultures were generated using these cell lines. In the endpoint, classification of cancer cells as MDR or non-MDR was performed by combining drug screening cell cytotoxicity assay, real-time monitoring of drug uptake and ROS, and flow cytometry-based confirmation of the MDR profile.

As reported previously^25,26^, biopsies were excised from the tumors of gingiva, tongue, and hypopharynx and harvested immediately for establishing monolayer cell lines *in vitro*. MCSs and 2D monolayer cells were developed using the same cells. Categorical segregation of MDR and non-MDR cancer cells was performed using a combination of anticancer drug screening on 2D and MCSs, differential uptake of calcein-AM in time-lapse fluorescence microscopy, monitoring of real-time ROS generation in the MCSs and 2D cultures, and a flow cytometry-based MDR assay. Finally, MCSs obtained from LK0917 gingiva tumor (referred to as MCS_17_ hereafter), LK0902 tongue tumor (referred to as MCS_02_ hereafter), and LK1108 hypopharynx tumor (referred to as MCS_08_ hereafter), were randomly selected for the development of a multidrug cancer resistance model system.

### Generation of MCSs from patient-derived HNSCC using forced floating method

The patient-derived HNSCC cell lines LK0917, LK0902, and LK1108 were revived from frozen stocks in 10 mL complete keratinocyte serum-free growth medium (KSFM, Gibco, Thermo Fisher Scientific), supplemented with 10% fetal bovine serum (FBS, Gibco), and penicillin 50 IU/mL and streptomycin 50 μg/mL (Thermo Fisher Scientific), and incubated in a humidified 5% CO_2_ atmosphere at a temperature of 37° C. Once cells reached 80% confluence, single cell suspensions were prepared by detaching the cells via mild enzymatic dissociation using 0.25% trypsin and 0.02% EDTA solution (Thermo Fisher Scientific). Trypsin was inactivated by adding complete KSFM medium. The number of live cells/mL were determined by adding 10 μL of 0.4% trypan blue (Thermo Fisher Scientific) to 10 μL of single cell suspension, mounting the mixture on Luna cell counter slides, and counting the cells on the automatic Luna cell counter. For the generation of MCSs sized 300–500 μm, 200 μL of LK0917 (MCS_17_), LK0902 (MCS_02_), and LK1108 (MCS_08_) single cell suspensions were seeded in ULA plates (Corning Life Sciences) at varying cell densities in the range of 0.25–0.75 × 10^5^ cells/mL. The plates were incubated at a humidified 5%CO_2_ atmosphere at 37°C (48-72 hrs) for maturation and assessment of MCSs diameter variation with respect to cell density. Progression of spheroid formation was imaged on a daily basis using a bright field microscope with 5× or 10× objectives and further image analysis was performed. Formation of MCSs was also monitored every 3 hours by live-cell imaging using Incucyte Zoom™ throughout the entire spheroid formation process with a phase-contrast set up using the 10× objective, and the images were analyzed.

### In vitro drug screening assay on 2D cell cultures and MCSs

Single cell suspensions of LK0917, LK0902, and LK1108 cell lines were seeded in 96-well flat bottom plates at a cell density of 8000 cells/well in 200 μL complete medium at 37°C and 5% CO_2_ atmosphere for 24 hours before drug treatment. After 24 hours, the culture medium was carefully aspirated and 2D cultures of three cell lines were treated with cisplatin (1, 2, 4, 6 & 8 μg/mL), doxorubicin (0.1, 0.2, 0.4, 0.6 & 0.8 μg/mL), and methotrexate (1, 2, 4, 6 & 8 μg/mL) prepared from their stock solutions (1 mg/mL) in complete KSFM medium. Cells were treated with drugs for 72 hours. Generation of MCSs was performed as described in the previous section. The cell density for the cytotoxicity assays was 0.7 × 10^5^ cells/mL for both MCS_17_ and MCS_02_ and 0.5 × 10^5^ cells/mL for MCS_08_. Tumor spheroids were incubated at 37°C and 5% CO_2_ atmosphere for 48 hours. After 48 hours of spheroidization, MCS_17_, MCS_02_, and MCS_08_ were treated with different doses of cisplatin, doxorubicin, and methotrexate at the same concentrations used for the 2D cell cultures, by replacing 50% of the culture medium with freshly prepared drug-supplemented^29^ medium, followed by incubation at 37°C and 5% CO_2_ atmosphere for 72 hours. For each drug concentration, 8 MCSs were used in triplicates, with effective drug concentrations equivalent to those used for the 2D cell cultures. Cell cytotoxicity in the drug-treated 2D cell cultures was assessed using the CellTiter96^®^ AQueous One Solution Cell Proliferation Assay (Promega). Briefly, at the end of 72 hours, the drug supplemented medium was replaced with 317 μg/mL MTS reagent-supplemented medium. For a total volume of 200 μL, 40 μL of the MTS reagent was added into each well and the plates were incubated at 37°C and 5% CO_2_ atmosphere for 3 hours. At the end of the incubation period, absorbances at 490 nm and 650 nm were recorded using a microplate reader (VersaMax™, Molecular Devices). All experiments were performed in triplicates.

### Real-time monitoring of calcein-AM uptake in 2D cell cultures and MCSs using fluorescence live-cell imaging

Acetoxymethyl ester (AM) derivatives of fluorescent probes such as calcein are actively pumped out of cancer cells with higher Pg-p and MRP1 expression^28^. In the present context, we have utilized the enhanced efflux properties of MDR tumor cells to generate separate calcein-AM uptake kinetic profiles for cell monolayers and MCSs. We monitored the real-time calcein uptake and intracellular calcein accumulation in 2D cell cultures and MCSs of LK0917, LK0902, and LK1108 using live-cell fluorescent imaging over a period of 11 hours (images represent up to 10 hrs) with image acquisition at 20-minute intervals.

In brief, cells were seeded in 96-flat bottomed plate (monolayers) and ULA (for MCSs) at a density of 0.7 × 10^5^ cells/mL (monolayers) and cultured for 24 hours before the start of the experiment. MCS_17_, MCS_02,_ and MCS_08_ were generated as described in the previous section. After spheroid formation, the KSFM growth medium was carefully decanted without disturbing the spheroids. Monolayer cell cultures and MCSs were incubated in serum-free KSFM medium containing non-fluorescent calcein-AM (1 mM in DMSO, Sigma) at a final concentration of 1 μM, for 12 hours at 37°C and 5% CO_2_ atmosphere. During the 12-hour incubation period, phase contrast and green fluorescence (Calcein_Ex/Em_ = 495/515 nm) images of the monolayer/spheroids were acquired every 15 minutes using time-lapse fluorescent microscopy. A 10× objective was used for image acquisition.

### Live-cell imaging of MCSs for calcein-AM uptake with varying cell density

For this experiment, MCS_17_, MCS_02,_ and MCS_08_ were prepared using various cell densities. Here, we seeded 1 × 10^4^, 1.5 × 10^4^, and 5 × 10^4^ cells/well for MCSs formation and assessed the calcein-AM uptake of the generated MCSs using the same procedure described in the earlier section.

### Monitoring of intracellular ROS generation in cell monolayers and MCSs using DCFDA assay and live-cell fluorescent microscopy

Monolayer cells were seeded at a density of 0.7 × 10^5^ cells/mL and cultured in complete KSFM medium for 24 hours before the experiment. MCS_17_, MCS_02_, and MCS_08_ were generated as previously described, and the complete growth medium was replaced with serum-free KSFM containing DCFDA (20 μM). The ULA plates containing the spheroids were immediately incubated in the Incucyte Zoom™ live-cell imaging microscope at 37°C and 5% CO_2_ atmosphere. Green fluorescence images were automatically obtained every 20 minutes for a total duration of 60 minutes.

### Flow cytometry-based assessment of specific MDR pump involvement

For the experiments on 2D cell cultures, single-cell suspensions of LK0917, LK0902, and LK1108 cell lines were prepared by trypsinization and counted using an automated cell counter as described previously. For each cell line, 1 × 10^6^ cells/mL were prepared in complete KSFM medium. For each sample to be assayed, 4 sets of tubes were prepared in triplicates.

For the experiments on MCSs, the spheroidization process for MCS_17_, MCS_02,_ and MCS_08_ was initiated 48 hours before performing the assay. After spheroidization, single cell suspensions were prepared from MCSs using 0.25% trypsin and 0.02% EDTA solution. The trypsinization time for MCS_17_ and MCS_02_ were 10 minutes and for MCS_08_ was 20 minutes. Immediately following trypsinization, complete KSFM medium was added in a 1:1 ratio. The MCSs were gently pipetted several times for complete dissociation. In the following step, different MDR pathway inhibitors such as novobiocin (BCRP inhibitor), verapamil (MDR1 inhibitor), and MK-571 (MRP inhibitor) provided with the MDR assay kit (Abcam 204534), were added to the reaction tubes. Complete KSFM medium containing 5% DMSO was used as a vehicle control. The reaction tubes were then incubated at 37°C for 5 minutes after gentle mixing, followed by the addition of efflux green detection reagent, gentle mixing, and incubation at 37°C for 30 minutes. Following 30 minutes of incubation, 5 μL of propidium iodide (PI) provided with the kit was added to the reaction mixture before performing flow cytometry. The cellular green fluorescence signal of efflux green detection reagent was measured using BD FACSARIA III in the PI-negative cell population using identical PMT voltage settings. Mean fluorescence intensity (MFI) values were calculated for each triplicate set of reaction tubes using the DIVA software and FlowJo 2.0.

### Microscopy image analysis

To obtain the automated real time drug uptake information, accurate segmentation of MCSs is important for quantitative analysis of red and green channel fluorescence of the spheroids. Intensity inhomogeneity over the spheroids and poor contrast in the boundary of spheroids are the major bottlenecks for acceptable segmentation. Traditional image segmentation techniques such as thresholding, region growing, and level set methods are unable to segment the spheroid with sufficient accuracy.

Therefore, we have used the P-Net (**Scheme 2a**) based fully convolutional network,^30^ which takes an entire image as input and returns a dense segmentation. The detailed architecture of P-Net is shown in **Scheme 2a**. The first 13 convolution layers of P-Net were grouped into five blocks, where the first and second blocks were each composed of two convolution layers, and each of the remaining blocks were composed of three convolution layers. The size of the convolution kernel was fixed as 3 × 3 in all convolution layers. Dilated convolution^31^ was used in P-Net to preserve the resolution of feature maps and enlarge the receptive field to incorporate larger contextual information. Images were resized to 512 × 512 pixels to reduce the time of segmentation. Several augmentation techniques such as flip and rotation were performed to increase number of training images. A stochastic gradient-based optimization ADAM^32^ was applied to minimize the cross-entropy based cost function.

**Scheme 2:**
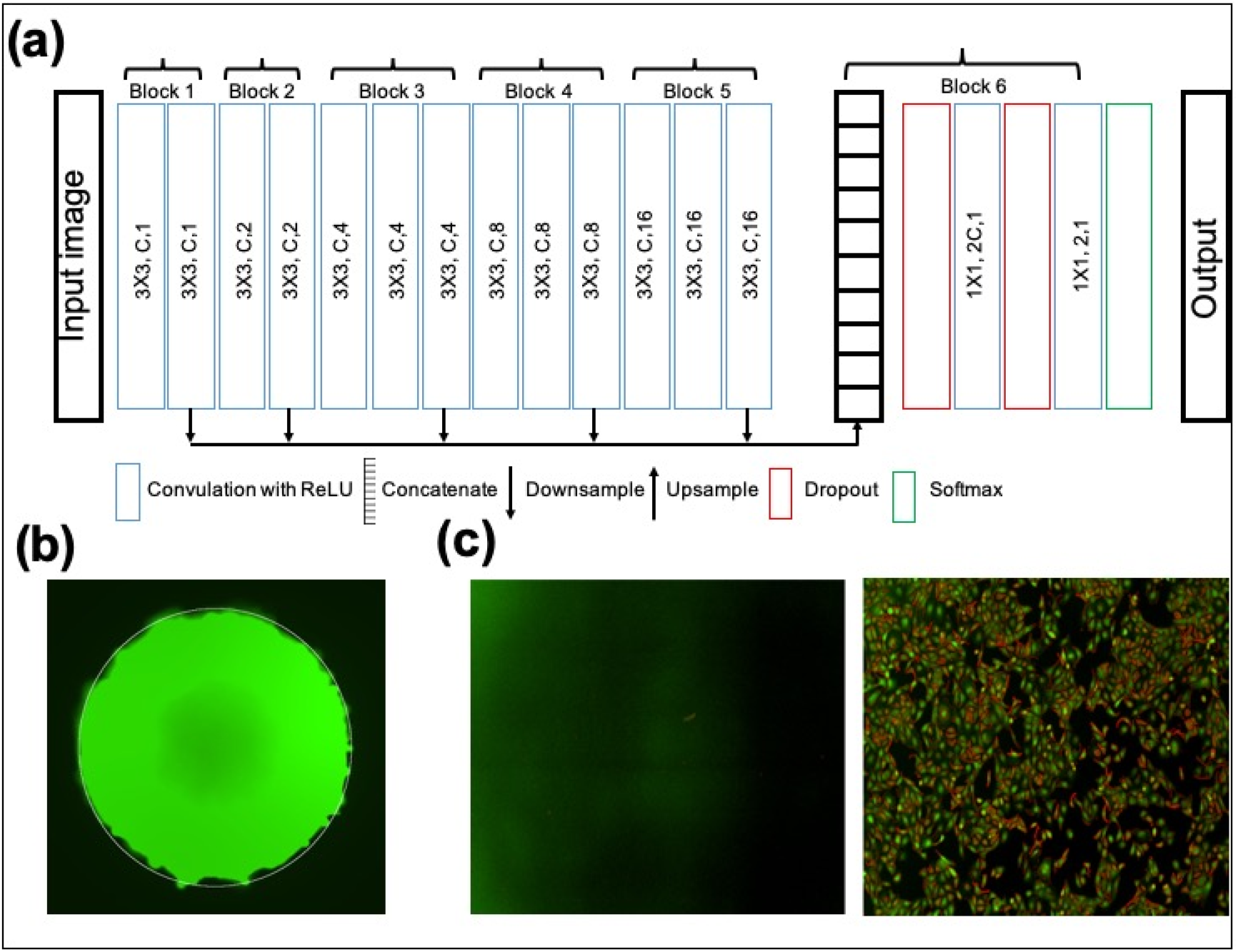
(a) Architecture of P-Net, (b) Segmentation of MCSs using P-Net and (c) control-without fluorescence (upper panel) and calcein-AM uptake by 2D cultures (lower panel).

The learning rate for the ADAM optimizer was set to 0.0001 and over-fitting was reduced by using dropout.^33^ The background and foreground weights were maintained at 1:10 ratio and training was performed up to 20 epochs.

The hyper-parameters were determined based on the validation dataset. The qualitative segmentation results are shown in **Scheme 2b**. Mean value of the green and red channels of the segmented spheroids indicate green fluorescence and red fluorescence, respectively. In the case of green fluorescence in 2D cell cultures, a Laplacian or Gaussian filter was applied to extract the edges of different pathological regions of the 2D cell culture images (**Scheme 2c**). The mean green fluorescence value over the edges of pathological regions of monolayer images was taken as a measure of green fluorescence.

### Statistical analysis

2D cell culture and MCS image analyses were performed using MATLAB 2016b (MathWorks, USA). ANOVA and Tukey’s multiple comparison test was performed in GraphPad Prism 8 for comparing different data sets. Values are presented as mean ± S.D. A p value < 0.05 was considered statistically significant. All experiments were performed in triplicates.

## Results

### Comparative drug response profiles of 2D cell cultures and MCSs

We studied the drug response profiles of 2D and MCSs obtained from LK0917, LK0902, and LK1108, to doxorubicin (**Fig. 1a**), cisplatin (**Fig. 1b**), and methotrexate (**Fig. 1c**), and compared drug efficacy and sensitivity between the two model systems (**Fig.1**).

**Figure 1.**
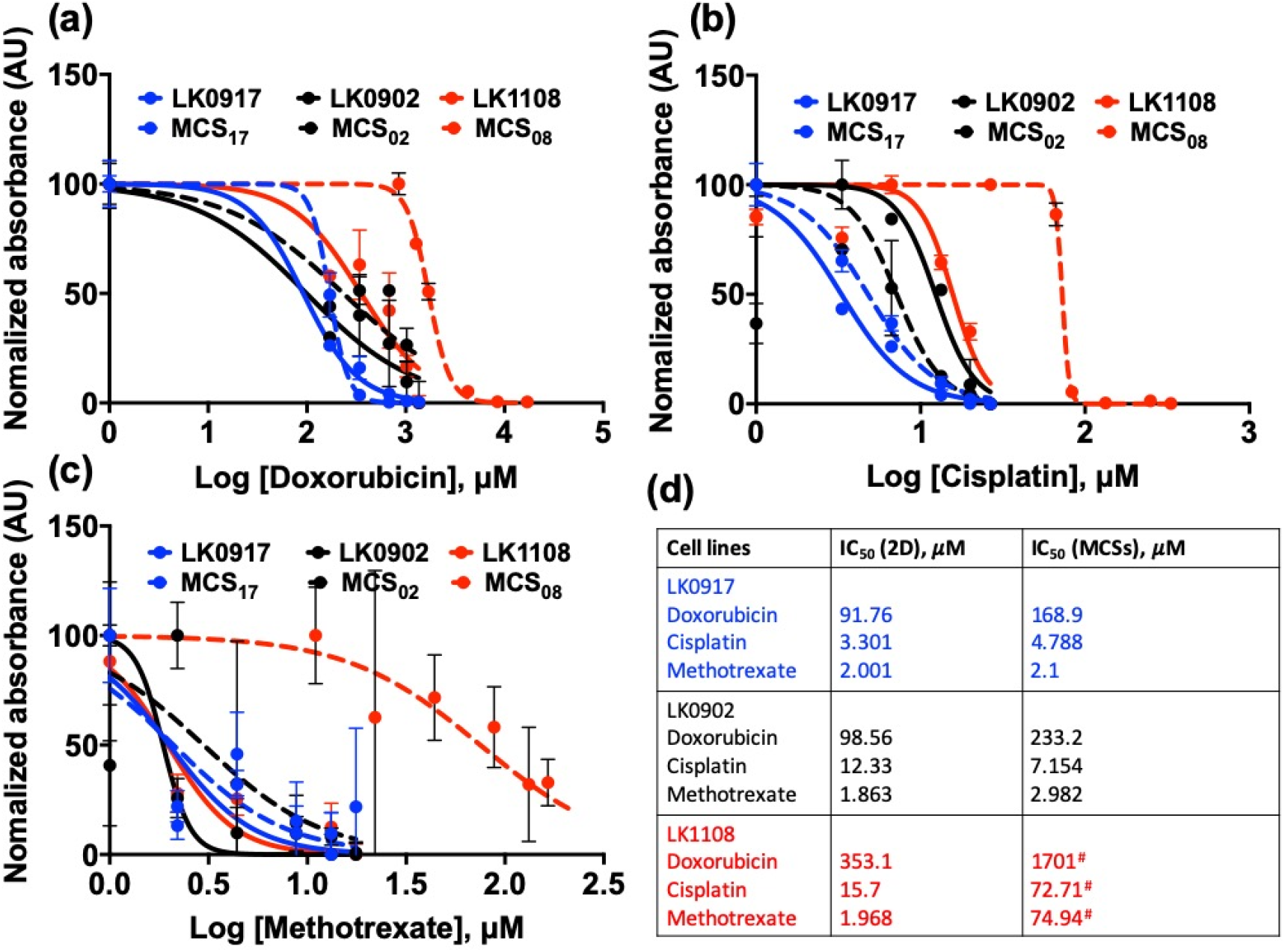
Comparative drug response profiles of 2D cell cultures and MCSs. (a-c) Drug response curves for LK0917, LK0902, and LK1108 for 2D and MCSs cultures treated with 0.1, 0.2, 0.4, 0.6, and 0.8 μg/mL of doxorubicin; 1, 2, 4, 6, and 8 μg/mL of cisplatin; and 1, 2, 4, 6, and 8 μg/mL of methotrexate. (d) IC_50_ (μM) values calculated from the drug response curves for both 2D and MCSs of all cell types for all drugs. Significantly large differences between the IC_50_ values of 2D and MCSs are denoted with “^#^”. Each IC_50_ value is the average of three independent experiments (n=3).

In our assessment, we included cisplatin and methotrexate, which are both drugs that are clinically approved for HNSCC treatment, with well documented activities^34,35^. In both the 2D cultures and MCSs of all cell lines, we observed lowest sensitivity to doxorubicin (lowest IC_50_ in comparison to the other two cell lines), followed by cisplatin and methotrexate. In terms of drug resistance, LK1108 appeared to be the least sensitive cell line to treatment having the highest IC_50_ values for all the three drugs tested, followed by LK0902, and LK0917. Interestingly, although large differences could not be observed between the drug responses of 2D cell cultures and MCSs obtained from LK0902 and LK0917 cell lines to cisplatin and methotrexate, significant differences were observed between the drug responses of LK1108 2D cell cultures and MCSs to doxorubicin. In addition, the difference in response of LK1108 2D cell cultures to cisplatin and methotrexate could not be observed in the LK1108 MCSs (**Fig. 1a-d**); however, we could still observe a stronger drug resistance pattern independent of the cell culture method used, thus providing an initial threshold of resistance pattern for the three HNSCC cell lines used in the present study.

### Real time monitoring of efflux pump activity in the 2D and MCSs of HNSCC cell lines using the calcein-AM uptake assay

We did not observe a significant difference in the calcein-AM uptake profiles of 2D cultures obtained from the LK0917, LK0902, and LK1108 cell lines, indicating that this *in vitro* monolayer model system might have limited use for the assessment of efflux pump activity, which is a direct measure of resistance (**Fig. 2a**).

**Figure 2:**
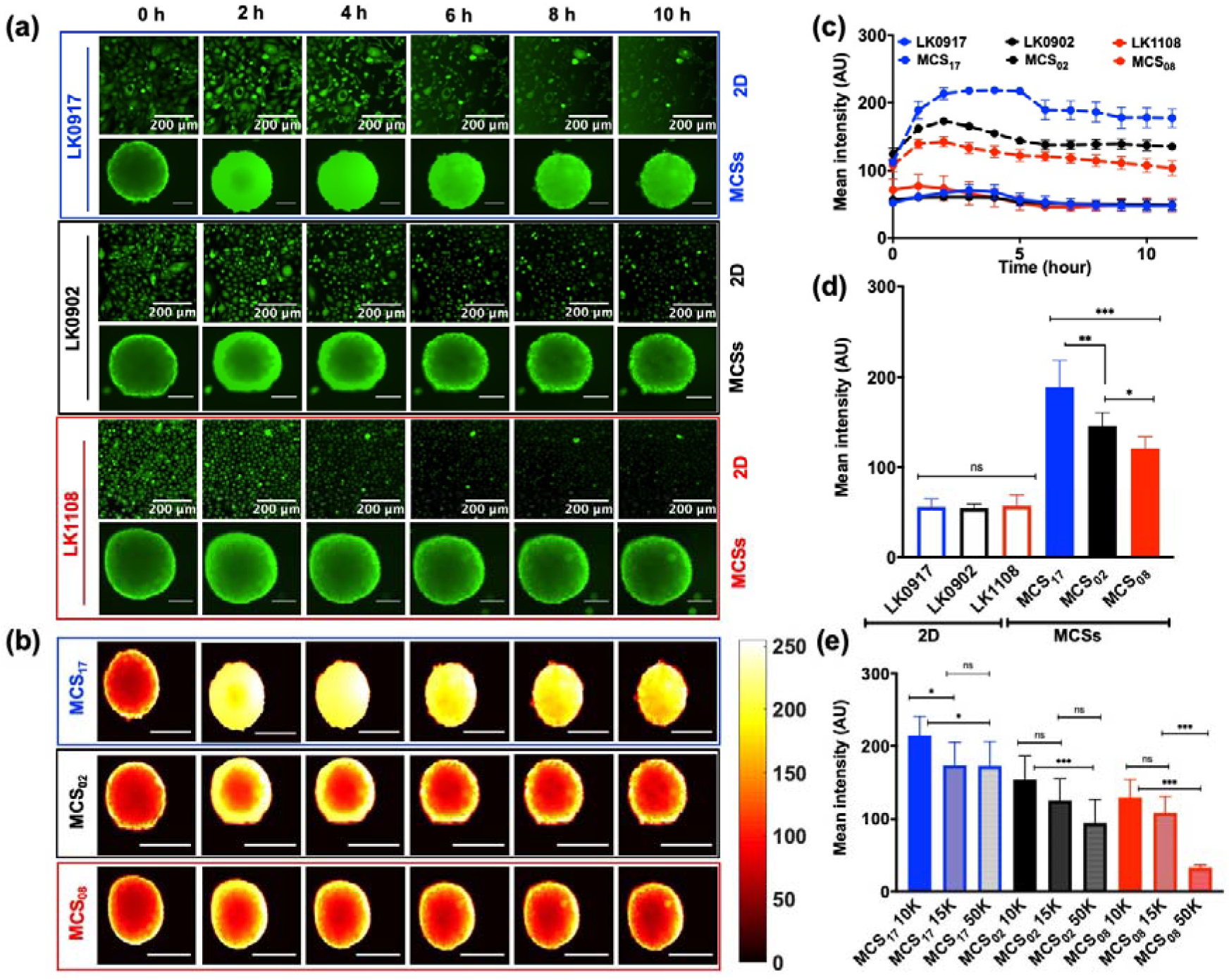
Real-time monitoring of calcein-AM uptake in 2D and MCSs. (a) Calcein-AM uptake in 2D cell cultures and MCSs obtained from LK0917-MCS_17_ (upper panel), LK0902-MCS_02_ (middle panel), and LK1108-MCS_08_ (lower panel) and over a time span of 10 hours with image acquisition at 20-minute intervals (scale bar, 200 μm). (b) Heat map pseudo color images of MCSs for differential calcein-AM uptake for the three cell lines (scale bar, 200 μm). (c) Mean fluorescence intensity profile with respect to time in the 2D cell cultures (LK0917, LK0902, and LK1108) and MCSs (MCS_17_, MCS_02_, and MCS_08_) respectively. (d) Total accumulated calcein over time in 2D and MCSs for all three cell lines. (e) Total accumulated calcein profiles of MCSs obtained from different cell densities. The data are shown as mean ± SD; ***p < 0.001, **p = 0.001, *p = 0.019, and ns = non-significant (n=3).

On the other hand, live cell fluorescent imaging showed significant differences between the calcein-AM uptake profiles of the MCSs generated from the three cell lines over time (**Fig. 2a** green fluorescence (upper, middle and lower panel) and **Fig. 2c**, mean fluorescence intensity over time). Pseudo-coloring mapping for the MCSs represented in **Fig. 2b**, exhibits the same calcein-AM uptake pattern for MCS_17_ (upper panel), MCS_02_ (middle panel), and MCS_08_ (lower panel) as exemplified in **Fig. 2a**. Maximum intra-spheroid green fluorescence, and thus maximum calcein retention, was observed for MCS_17_ followed by MCS_02_ and MCS_08_, which suggested that efflux pump activity was lowest in MCS_17_ followed by MCS_02_ and MCS_08_, consistent with the drug response profiles provided in **Fig. 1**. MCS_08_ was the most resistant to drug treatment, indicated by significantly higher IC_50_ values (**Fig. 1d**) compared to those of MCSs from other cell lines and indeed, the MCSs from these cells showed the highest efflux of calcein-AM over time, indicated by lowest green fluorescence over time, without penetration to the spheroid core. Likewise, MCS_17_ was the least resistant to drug treatment, indicated by significantly lower IC_50_ (**Fig. 1d**) values compared to those of MCSs from the other cell lines, which showed lowest efflux of calcein-AM over time, indicated by the highest green fluorescence over time, with penetration into the spheroid core and without complete expulsion over a period of 10 hrs. These findings suggest that MCS_17_, MCS_02_, and MCS_08_ can be characterized as treatment sensitive, moderately resistant, and highly resistant, respectively, based on their efflux pump activity (**Fig. 2a**). Not surprisingly, we observed a clear distinction between the mean fluorescence intensity over time for MCSs obtained from all three cell lines, while such a distinction could not be made for the 2D cultures of these cell lines (**Fig. 2d**). Consistent with the fluorescence profiles over time, the mean green fluorescence was lowest for MCS_08_ and highest for MCS_17_, indicating low and high efflux pump activities, respectively.

During the initial spheroidization process, we observed that MCSs obtained from different cancer cell lines had different spheroid diameters despite the seeding density for all cell lines was kept constant (**Supplementary section, Fig. S1**). In order to eliminate the possibility that spheroid size affected calcein-AM uptake, we performed a differential calcein uptake study using varying cell seeding densities for the initiation of spheroidization and found that calcein-AM uptake was independent of spheroid diameter (**Fig. 2e**). Interestingly, increasing seeding density appeared to be associated with decreased mean green fluorescence for the cell lines termed moderately and highly resistant, while for the treatment sensitive cell line, this pattern was not observed at the highest seeding density (**Fig. 2e**).

### ROS activity and MDR profiles of 2D cell cultures and MCSs obtained from HNSCC cell lines

Bidirectional modulation of ROS activity has been reported to induce MDR^38^. We used a ROS activity assay based on the same principle as the calcein-AM uptake assay, where DCFDA, which once intracellularly incorporated, first becomes deacetylated by cellular esterases into a non-fluorescent form, and then becomes converted to a highly fluorescent hydrophilic form that is retained in the cytosol upon oxidation by ROS.^36^ It is thus expected that increased intracellular and intra-spheroid green fluorescence would indicate ROS activity. While we could not detect a significant difference between the fluorescence of 2D cell cultures (**Fig. 3a**), fluorescence of MCSs differed significantly between different cell lines (**Fig. 3a and c**). We observed decreased green fluorescence over time in the MCSs obtained from the drug sensitive LK0917 cell line (**Fig. 3a**, upper panel), which indicates the presence of ROS activity that subsides over time in the absence of treatment. The lack of fluorescence in the moderately and highly resistant MCS_02_ and MCS_08_ cell lines (**Fig. 3a and c** middle and lower panel), respectively, indicated lack of ROS activity in the untreated MCSs. Subsequently, mean fluorescence intensity for both cultures were estimated over a time span of 60 minutes (**Fig. 3b**), indicating a distinct pattern for MCSs, whereas no significant differences were observed for the 2D cultures.

**Figure 3.**
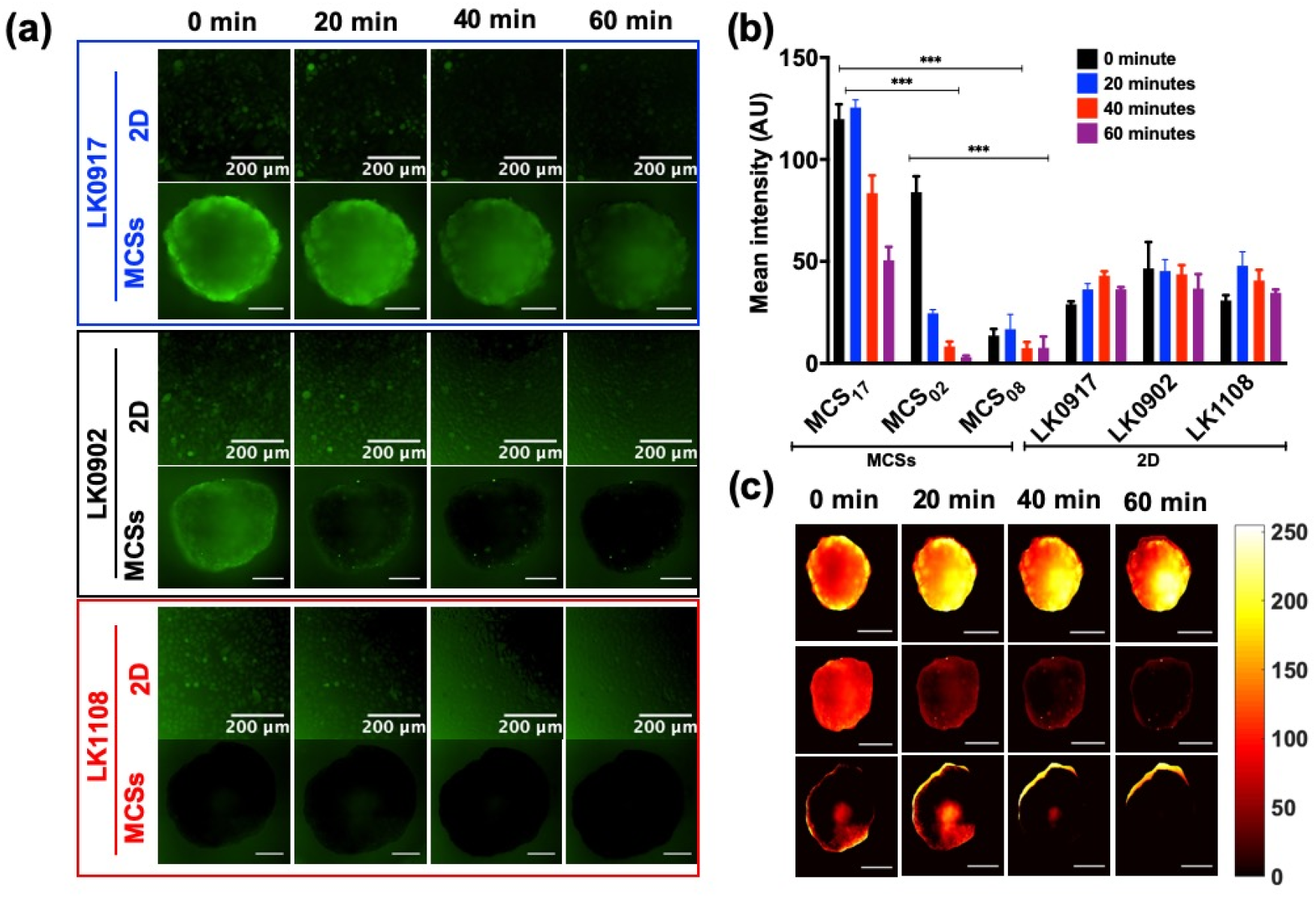
ROS activity of all cell lines in 2D and MCSs. (a) Live-cell fluorescence images of the 2D (LK0917, LK0902, and LK1108) and MCSs (MCS_17_, MCS_02_, and MCS_08_) cultures obtained from cell lines in the presence of DCFDA over a period of 60 minutes with image acquisition at 20-minute intervals (upper, middle and lower panel respectively), scale bar, 200 μm). (b) Redox state in 2D and MCSs cultures. Data are presented as the mean ± SD; ***p < 0.001, **p = 0.001, *p = 0.019, and no significant differences were observed in case of 2D (n=3). (c) Heat map pseudo color images of MCSs for differential redox status for the three cell lines (scale bar, 200 μm).

### Flow cytometry based MDR assay for the characterization of efflux pump activity in the HNSCC cell lines

P-gp, MRP1, and BRCP transporter activities were assessed flow cytometrically by measuring the efficacy of selective inhibitors of these transporters in preventing the efflux of the efflux green detection reagent, which is a substrate for all three transporters.

We determined the median fluorescence intensity (MFI) values for the 2D cell cultures and MCSs obtained from LK0917, LK0902, and LK1108, in the presence and absence of the specific efflux pump inhibitors verapamil, novobiocin, and MK-571 against P-gp, BCRP, and MRP1, respectively, using flow cytometry analysis^37–39^. The MFI values for all transporters were comparable for both 2D cell cultures and MCSs as shown in **Fig. 4a-c**. LK0917, which we previously identified to be the most sensitive cell line among the three, exhibited a greater change in fluorescence intensity after inhibitor treatment. Highest retention compared to non-inhibitor treated cells was observed for the BCRP transporter, followed by the MRP1, and P-gp transporters. For the LK1108 cell line, which was previously identified to be the most resistant among the three, highest retention compared to the non-inhibitor treated cells was also observed for the BCRP transporter, followed by Pg-p, and the MRP transporters. On the contrary, for the moderately resistant LK0902 cell line, highest retention compared to non-inhibitor treated cells was observed for the P-gp and MRP1 transporters, followed by the BCRP transporter. The fact that the lowest MFI was observed for the most resistant cell line indicates overexpression of these transporters, thus suggesting ineffective inhibition of efflux pump activity. Likewise, in the cell line identified to be the most sensitive to treatment among the three, efflux pump activity was more effectively inhibited owing to lower efflux pump expression, indicated by a higher MFI value. Simultaneously, real-time fluorescence imaging was performed with one of the pump inhibitors (verapamil, shown in **Fig. 4e and f**).

**Figure 4:**
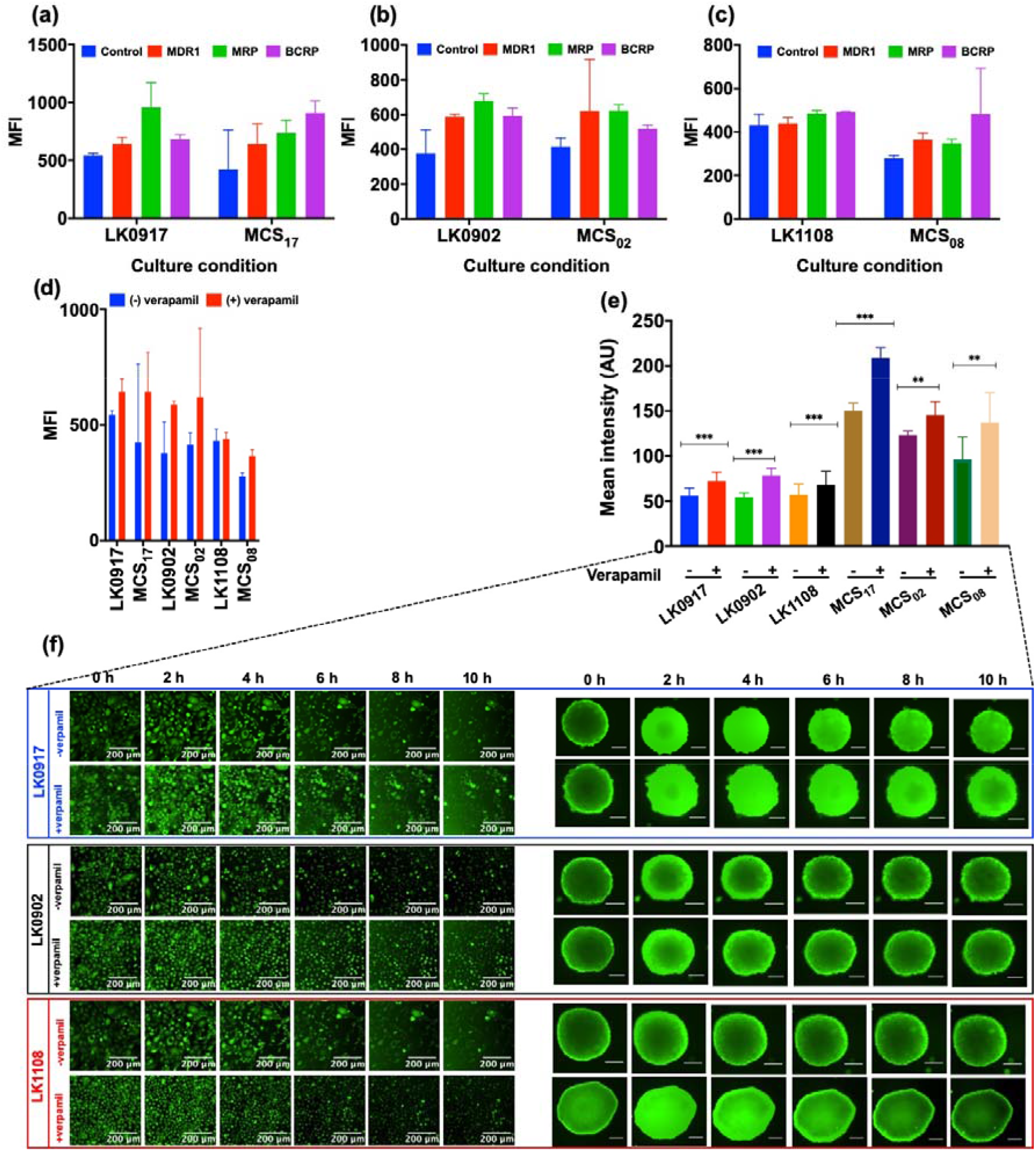
(a), (b) and (c) Median fluorescence intensity (MFI) values for 2D and MCSs cultures of LK0917, LK0902, and LK1108 in presence of different ABC pump inhibitors for MDR1, MRP, and BCRP. (d) MFI for MDR1 pump inhibitor (verapamil) and its corresponding live cell imaging for 2D and MCSs cultures. (e) Total calcein-AM uptake for the entire course of 10 hours +/− verapamil for 2D and MCSs. (f) elucidate calcein-AM uptake in 2D for LK0917, LK0902 and LK1108 in presence and absence of verapamil over a time span of 10 hours with images taken after every 20 minutes (images shown here are at 2 h interval). scale bar 200 μm. (c) The data are shown as a mean of ± SD, ***p<0.001, **p=0.001.

## Discussion

The global annual occurrence of head and neck cancers exceeds 0.5 million^40^, out of which 90% are HNSCCs. Early-stage disease progression is curable by surgical removal of tumor tissue and radiotherapy, but the prognosis for recurrent disease onset is still challenging and puzzling^41^. Presently, chemotherapeutic drugs such as cisplatin, 5-fluorouracil (5-FU), and taxanes such as paclitaxel and docetaxel, are the standard treatment options for recurrent or advanced HNSCC. However, the variability and robustness of these treatment modalities are not very well understood^42,43^. MDR against cytotoxic drugs is regarded as the main clinical impediment in using chemotherapy for the treatment of HNSCC. MDR is a result of the interplay between a diversity of factors, which include overexpression of the transporter molecules Pg-1, MRP1, and BCRP^44–50^. In spite of this well-known phenomenon, effective detection methods are still lacking for correct characterization of MDR status in cancer cells. Development of strategies that enable this characterization can prove to be highly effective in devising targeted treatment regimens against sensitive and resistant cancer cells. Presently, commonly used detection methods include polymerase chain reaction (PCR), in-situ hybridization (ISH), and RNase protection assays (RPAs) for the quantification of Pg-1 mRNA levels. Western blotting and immunohistochemistry have also been used for the detection of MDR proteins^51^. In the present study, we have attempted to simplify the identification of MDR status in patient-derived HNSCCs by combining drug screening, measurement of the difference in calcein-AM uptake studied using live-cell fluorescence imaging, fluorescence-based assessment of ROS activity, and flow cytometry-based prediction of ABC transporter involvement. We performed comparative assessment of 2D cell cultures and MCSs and observed noticeable differences between the two *in vitro* systems. 3D tumor spheroids were introduced as model systems by Sutherland et al., owing to their resemblance to solid tumors in many structural and microenvironmental aspects, and they serve as the most reliable *in vitro* model for investigating therapeutic and mechanistic approaches^52^.

In the present study, we have used 2D cell cultures and MCSs obtained from the patient-derived LK0917, LK0902, and LK1108 HNSCC cell lines. Firstly, we performed treatment response cell viability assay in the presence of cisplatin, doxorubicin, and methotrexate to assess drug cytotoxicity in both 2D cell cultures and MCSs (Fig. 1). Overall, cells grown as MCSs showed higher IC_50_ values compared to the 2D cultures. Interestingly, LK1108 cells required the highest dose for all the three drugs in order to achieve 50% inhibition **(Fig. 1d)**, irrespective of the culturing method, which indicated that among the three cell lines, LK1108 showed the highest resistance to treatment. On the other hand, lowest IC_50_ values were observed for LK0917, indicating this cell line as the most drug sensitive among the three cell lines. These findings suggested MDR status of LK0917<LK0902<LK1108.

Next, we performed live-cell fluorescence imaging of calcein-AM uptake in the 2D cell cultures and MCSs, in order to assess transporter activity, as calcein-AM is a substrate of P-gp and MRP1^53–56^. We have chosen live-cell fluorescence imaging because dynamic cellular changes can be observed using this method unlike fixed-cell imaging. In addition, real-time fluorescence microscopy is less prone to experimental artifacts, rendering the outcomes more reliable. We did not observe any difference between the calcein-AM uptake profiles of 2D cell cultures obtained from LK0917, LK0902, and LK1108 (**Fig. 2a, c**), with total accumulated calcein trapped inside the cells remaining comparable for the three cell lines, hence making it extremely problematic to detect differences in drug resistance activity of these cell lines (**Fig. 2c, d**). On the contrary, using MCSs as the *in vitro* model system, we were able to detect differences in drug resistance, indicated by calcein accumulation over time. Over a 12h time window, we observed highest calcein accumulation in MCS_17_, followed by MCS_02_ and MCS_08_, which indicated that transporter activity was lowest in the MCS_17_, followed by MCS_02_ and MCS_08_ (**Fig. 2a and c**). Comparison of the mean green fluorescence intensities between the 2D cell cultures and MCSs confirmed these findings (**Fig. 2d**). In order to assess whether spheroid diameter affected calcein-AM diffusion, we performed the same experiments with varying cell seeding densities and observed the same calcein accumulation pattern, indicating that spheroid diameter did not have any effect on the calcein-AM uptake profiles of MCSs, thus eliminating the possibility of false positives (**Fig. 2e**). After establishing the transporter activity based MDR status in these cell lines, we further validated our findings using the DCFDA assay for the detection of ROS activity.

DCFDA is a hydrophobic fluorogenic dye used in the measurement of intracellular ROS activity. After cellular uptake, DCFDA is deacetylated by cellular esterases into a non-fluorescent hydrophilic compound that cannot exit cells, which is further oxidized into a highly fluorogenic product in the presence of ROS, which can be detected by fluorescence microscopy. We observed highest ROS activity in MCS_17_, indicated by green fluorescence and no ROS activity indicated by a lack of fluorescence in the MCS_02_ and MCS_08_ spheroids (**Fig. 3a-c**) that we identified to have higher drug resistance compared to the MCS_17_. ROS activity is indicative of oxidative stress. It is expected that drug resistant cell lines that actively pump drugs out of the cell, have acquired higher survival capacity compared to those that cannot. In this context, lack of ROS activity in the untreated MCS_02_ and MCS_08_ spheroids, indicates lack of ROS activity in these cells, which could be attributed to another mechanism of survival, as oxidative stress is detrimental to cell survival, thus indirectly supporting our finding that these cell lines are highly drug resistant, one of the mechanisms being high transporter activity and the other being acquired lack of ROS activity. On the other hand, the presence of ROS activity in the non-resistant MCS_17_ spheroids show that these cells are not in good condition owing to oxidative stress and therefore they are more responsive to drug treatment. These findings indicate that by assessing drug transporter activity and ROS activity, the MDR status of patient cancer cells can be further characterized based on their drug transporter and ROS activities, which could potentially help determine patients that can benefit from a particular treatment regimen.

## Conclusion

We have established an assay system for the determination of cancer cell MDR status in cancers with unknown drug resistance profiles. Using our assay system, we were able to predict the efflux pump activities of three different patient-derived HNSCCs, which is important for determining cytotoxic drug vulnerability and the potential of developing MDR as a result of repeated drug exposure. The methods we described here could potentially be integrated into translational research for obtaining the MDR status of cancers, and aiding in the determination of the optimal treatment strategy.

## Acknowledgment

MA and HP acknowledge funding from MIIC, PDF grant and seed grant from Linköping University, Sweden. HP acknowledge EU H2020 Marie Sklodowska- Curie Individual Fellowship (Grant no. 706694) from European Commission and Wolfson College, University of Cambridge for supporting HP with Junior Research Fellowship (B1). MA acknowledges IKE-LiU for core facility and laboratory set up to perform all the experiments under supervision of JH and KR. KR acknowledge the The Swedish Cancer Society (2017/301), the County Council of Östergötland, and the Research Funds of Linköping University Hospital.

## Notes

#### Summary of Updates

This version of the manuscript has been updated for recently revised figures and text.

