## Supplementary for "Multicellular tumor spheroids- an effective *in vitro* model for understanding drug resistance in head and neck cancer"

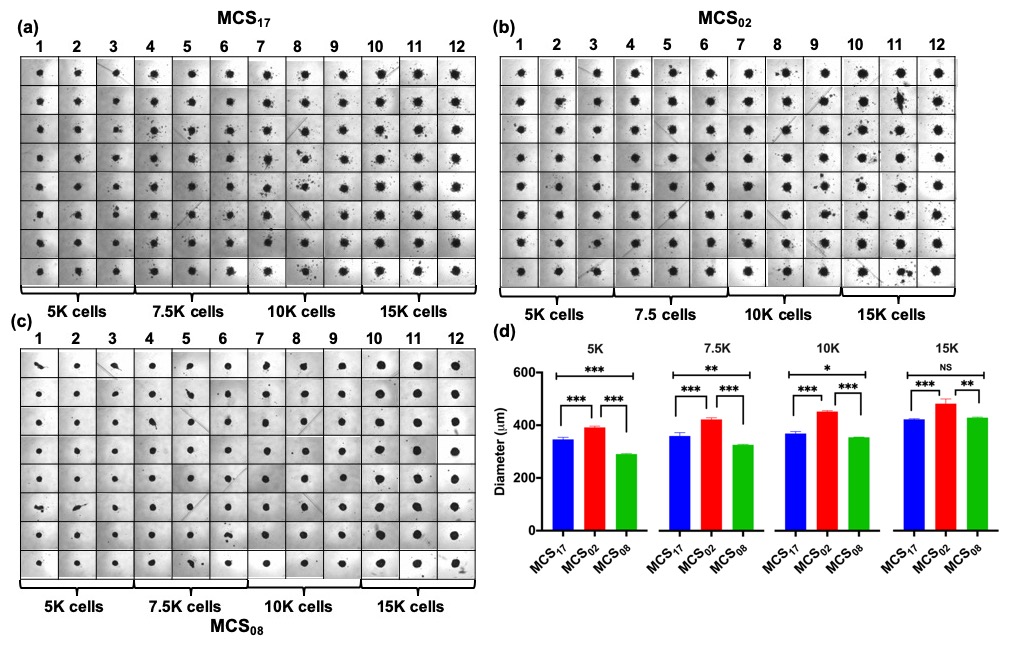

***Figure S1:*** *(a) – (c) Bright field images of MCSs (MCS_17_, MCS_02_, MCS_08_) for the three cell lines seeded at different cell density in ultra-low attachment plates acquired at 5x magnification. Cells were seeded at 5K, 7.5K, 10K, and 15K cells/ well and the corresponding diameter of the spheroids (b). The data are shown as a mean of ± SD, ***p<0.001, **p=0.001, *p=0.019.*
